# Comparison of single-cell whole-genome amplification strategies

**DOI:** 10.1101/443754

**Authors:** Nuria Estévez-Gómez, Tamara Prieto, Amy Guillaumet-Adkins, Holger Heyn, Sonia Prado-López, David Posada

**Affiliations:** Department of Biochemistry, Genetics and Immunology, University of Vigo, 36310 Vigo, Spain.; Biomedical Research Center (CINBIO), University of Vigo, 36310 Vigo, Spain.; Galicia Sur Health Research Institute, 36312 Vigo, Spain.; CNAG-CRG, Centre for Genomic Regulation (CRG), Barcelona Institute of Science and Technology (BIST), Barcelona, Spain; Universitat Pompeu Fabra (UPF), Barcelona, Spain

## Abstract

Single-cell genomics is an alluring area that holds the potential to change the way we understand cell populations. Due to the small amount of DNA within a single cell, whole-genome amplification becomes a mandatory step in many single-cell applications. Unfortunately, single-cell whole-genome amplification (scWGA) strategies suffer from several technical biases that complicate the posterior interpretation of the data. Here we compared the performance of six different scWGA methods (GenomiPhi, REPLIg, TruePrime, Ampli1, MALBAC, and PicoPLEX) after amplifying and low-pass sequencing the complete genome of 230 healthy/tumoral human cells. Overall, REPLIg outperformed competing methods regarding DNA yield, amplicon size, amplification breadth, amplification uniformity –being the only method with a random amplification bias–, and false single-nucleotide variant calls. On the other hand, non-MDA methods, and in particular Ampli1, showed less allelic imbalance and ADO, more reliable copy-number profiles and less chimeric amplicons. While no single scWGA method showed optimal performance for every aspect, they clearly have distinct advantages. Our results provide a convenient guide for selecting a scWGA method depending on the question of interest while revealing relevant weaknesses that should be considered during the analysis and interpretation of single-cell sequencing data.

---

Advances in single-cell genomics have made possible the study of genomic variation at the most basic level, rapidly generating many new insights into complex biological systems, from microbial diversity to immune response, development or tumor progression^1^. While single-cell RNA sequencing is now mature and almost standard, single-cell DNA sequencing is still quite challenging^2^, mainly due to an amplification step needed before the characterization of the genome, as it is not possible to directly sequence the 6-7 pg of DNA present, for example, in a human cell. While whole-genome single-cell library preparation without preamplification is possible^3^, these types of techniques still include several PCR cycles, usually rely on custom-made microfluidic devices, and their implementation in a standard laboratory is far from trivial. Therefore, single-cell whole-genome amplification (scWGA) is still a prerequisite in many applications of single-cell genomics.

Multiple scWGA methods have been proposed, typically based on pure PCR^4–7^, multiple displacement amplification (MDA)^8,9^ or a combination of both^10,11^, but always relying on the use of DNA polymerases. Unfortunately, the latter have a limited strand extension rate and processivity, and during scWGA lots of priming and extension reactions are required^12^. This large amount of reactions entails significant technical errors such as (1) allelic imbalance (AI) or allelic dropout (ADO) –when a particular allele is preferentially amplified or not amplified at all, respectively–, (2) non-uniform coverage usually attributed to GC content affecting denaturation and primer binding efficiency^13–15^, generation of chimeric DNA molecules due to the polymerase strand displacement activity^16–18^ and false single-nucleotide variants (SNVs) owing to the infidelity of the DNA polymerase^14^ (Fig. 1).

**Fig. 1.**
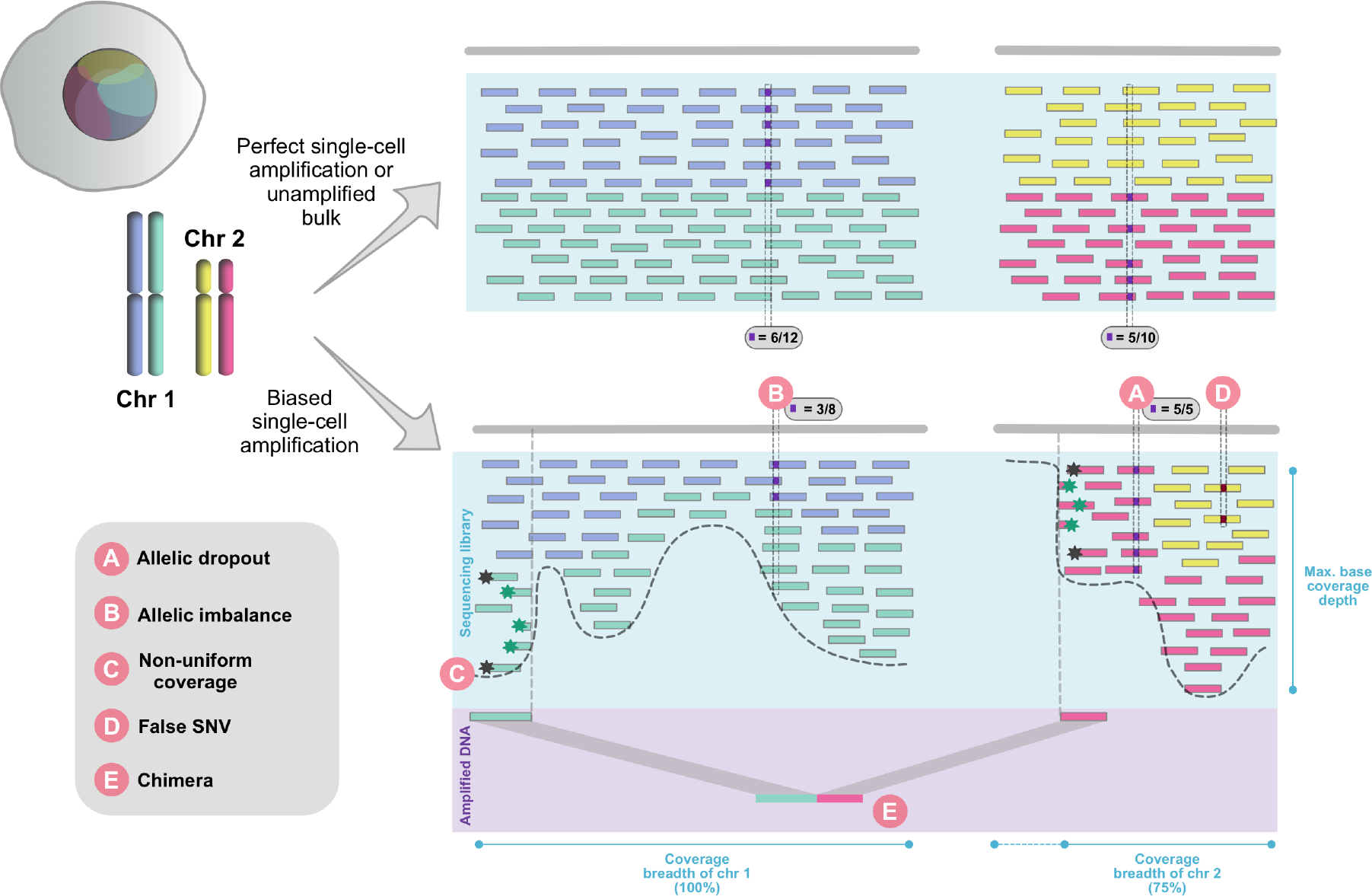
scWGA technical errors. On top, there is a representation of reads from a sequencing library constructed from perfectly amplified single-cell DNA material or, commonly, unamplified bulk. Reads coming from a sequencing library built from the product of a biased amplification are shown at the bottom. Single-cell amplification starts from a single pair of homologous chromosomes (Chr 1 and 2) while the unamplified bulk library is directly constructed from lots of chromosome pairs. During the amplification, which originated the second sequencing library, several biases occurred. On the one hand, some templates were not amplified at all (A: allelic dropout), or they were not copied as many times as their homologous sequences leading to a disproportion of maternal and paternal alleles (B: allelic imbalance). The latter contributes (although is not strictly required) to the coverage non-uniformity across the genome (C: Non-uniform coverage). The uneven coverage results in a decrease of the coverage breadth (proportion of genome covered by at least 1 read). On the other hand, during the amplification the DNA polymerase introduced a single-nucleotide variant not present in the original template (D: false SNV) as well as one chimeric amplicon (E: chimera), due to a replication error and a strand displacement, respectively. Discordant paired-end reads (grey stars) and split reads (green stars) reveal the presence of such chimeric amplicons. Little squares represent alternate alleles at original true germline sites (A,B) and one false SNV (D). Although amplicons are usually longer than reads, here they have been shortened to facilitate the representation.

While several studies comparing the relative performance of different scWGA strategies have already been published, their scope is usually limited in terms of the sequencing target, number of scWGA methods evaluated and/or number and type of amplified cells^16,17,19–30^ (Supplementary Table 1). To date, we are not aware of any study comparing a large number of scWGA strategies on whole genomes obtained from a large number of individual cells. Here we report a comprehensive benchmark of six popular scWGA kits, including five next-generation sequencing (NGS) library preparation kits and two NGS technologies, using three human cell lines. In total, we obtained 230 single-cell whole-genome sequences under 54 different scenarios (Fig. 2). We show that MDA and non-MDA methods perform differently for distinct purposes, and identify important differences within these categories. Our results should help single-cell genomics researchers choose the best amplification method for their question of interest.

**Fig. 2.**
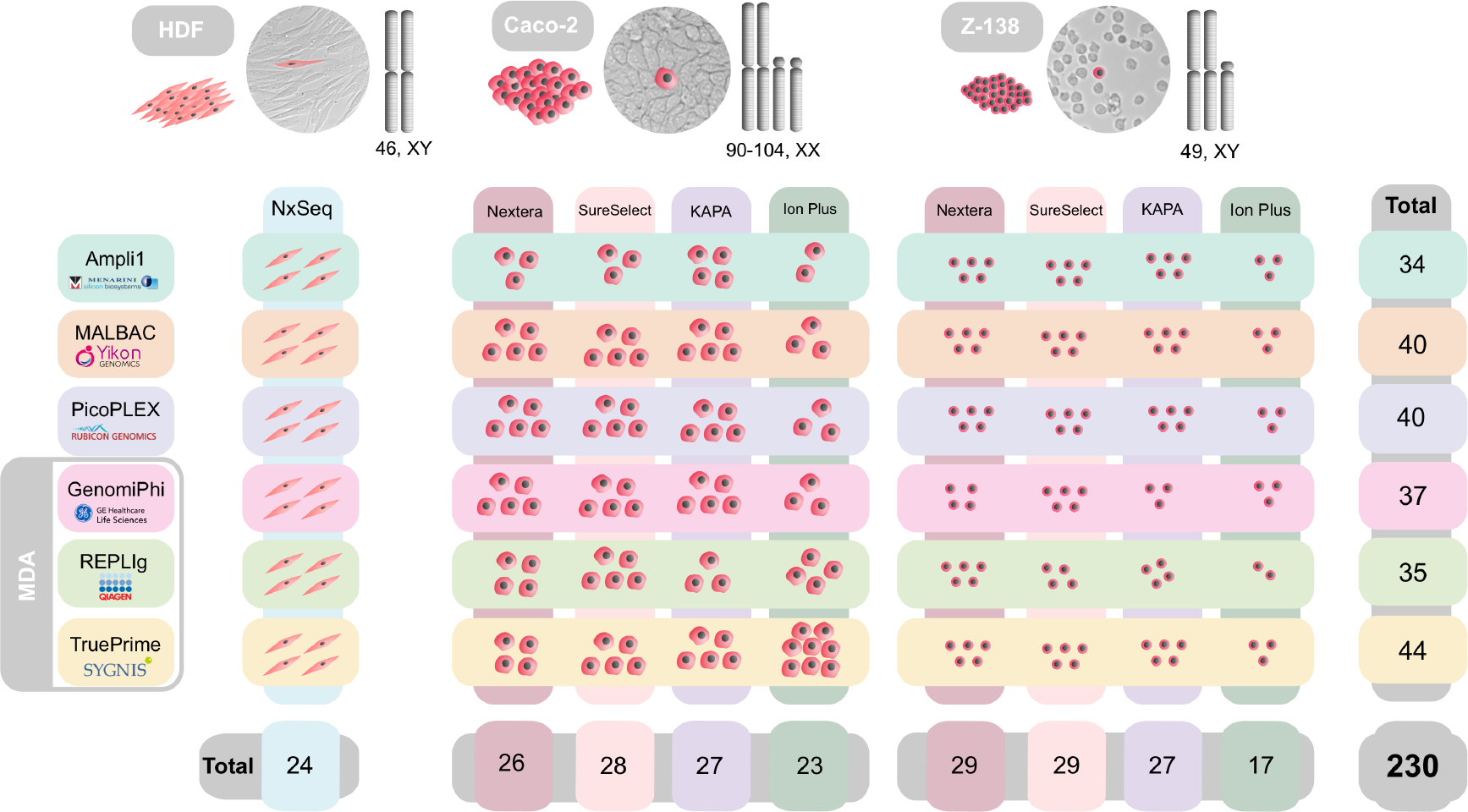
Experimental design. 24 HDF single-cells, 104 Caco-2 single-cells and 102 Z-138 single-cells sorted by FACS were amplified using six scWGA methods (three MDA and three non-MDA). Sequencing libraries were constructed following four different protocols for Caco-2 and Z-138 and one for HDF. A bulk sample from the HDF cell line was also extracted and sequenced as unamplified control.

## Results

We assessed the performance of six scWGA commercial kits, three MDA (GenomiPhi, REPLIg and TruePrime) and three non-MDA (Ampli1, MALBAC and PicoPLEX) (Supplementary Table 2), and five library preparation kits (Supplementary Table 3) in terms of amplification yield, amplicon size, coverage breadth, coverage uniformity, chimera formation, copy-number detection, allelic imbalance, ADO and false SNV detection. To do this, we obtained low-pass (0.07-1.8X) whole-genome sequencing (WGS) data from 230 individual human cells from a healthy fibroblast cell line (HDF), a colorectal cancer cell line (Caco-2), and a mantle lymphoma cell line (Z-138).

### DNA yield, amplicon size, and integrity

The amount of DNA obtained with the different scWGA kits, plus the size and quality of the amplicons, could be a limitation for downstream experiments. Here, we observed statistically significant differences among the scWGA methods for DNA yield and amplicon integrity and size, independently of the cell line (Fig. 3 and Supplementary Tables 4-6). REPLIg resulted in the highest DNA yield by far with a mean value across cell lines close to 35 μg, while the other scWGA kits produced average yields below 8 μg (Fig. 3a and Supplementary Tables 4-6). MDA approaches (GenomiPhi, REPLIg, TruePrime) produced amplicon sizes much larger than non-MDA methods (Ampli1, MALBAC, PicoPLEX) (around 10 and 1.2 kb on average, respectively), with REPLIg producing the largest amplicons (>30 kb) (Fig. 3c and Supplementary Tables 4-6). The size of the amplicons produced by the MDA kits allowed us to estimate amplicon integrity using the DNA Integrity Number (DIN) value, with REPLIg showing significantly higher values than GenomiPhi or TruePrime (Fig. 3b and Supplementary Tables 5,6). For all these three parameters, MDA methods were much more variable than non-MDA approaches, in particular in the case of REPLIg, which displayed the largest standard deviations (Supplementary Tables 4-6).

**Fig. 3.**
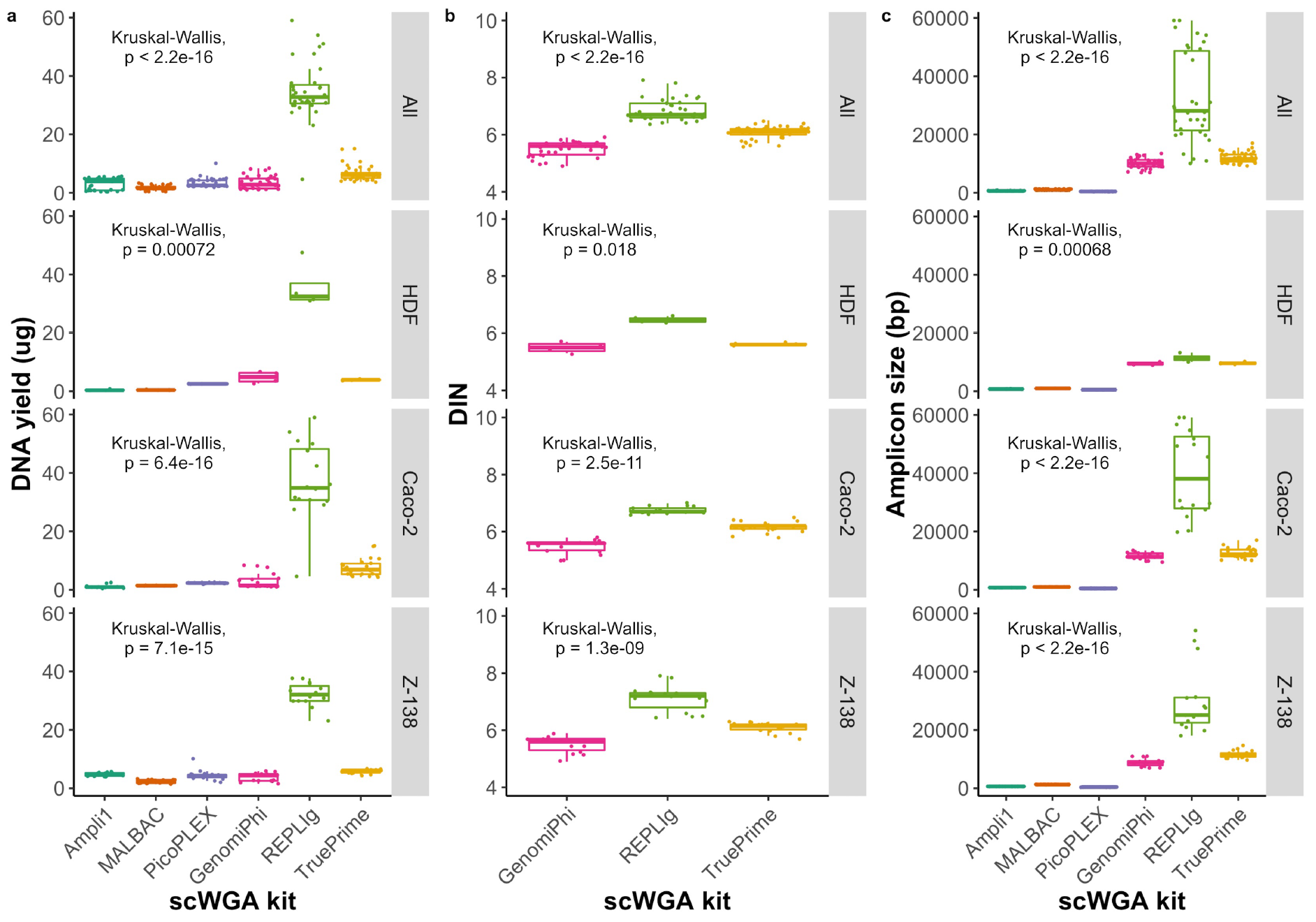
Amplification yield. **a,** DNA yield. **b,** DIN values (only MDA methods were measured). **c,** Amplicon size. Results are also shown combining the three cell lines (All). **a-c,** Boxplots: the central line indicates the median, while the box limits correspond to the Q1 and Q3 quartiles; upper and lower whiskers extend from Q3 to Q3 + 1.5 × (Q3 - Q1) and from Q1 to Q1 - 1.5 × (Q3-Q1), respectively. Kruskal-Wallis test p-values are shown for each dataset.

### Amplification breadth and uniformity

An ideal scWGA method should provide a set of DNA molecules that represent the target genome as completely as possible. If the amplification is not uniform, different genomic regions may be missed. Here, we observed statistically significant differences among scWGA methods for amplification breadth and uniformity for the different cell lines (Fig. 4a,b). Overall, REPLIg yielded the highest amplification breadth (∼50%) and TruePrime the lowest (∼15%) (Fig. 4a and Supplementary Tables 4-6). In general, REPLIg also resulted in a more uniform amplification –measured by the Gini index of the Lorenz curves– than the other scWGA methods, with TruePrime performing the worst (Fig. 4b; Supplementary Fig. 1 and Supplementary Tables 4-6). Again, MDA methods were much more variable than non-MDA approaches.

**Fig. 4.**
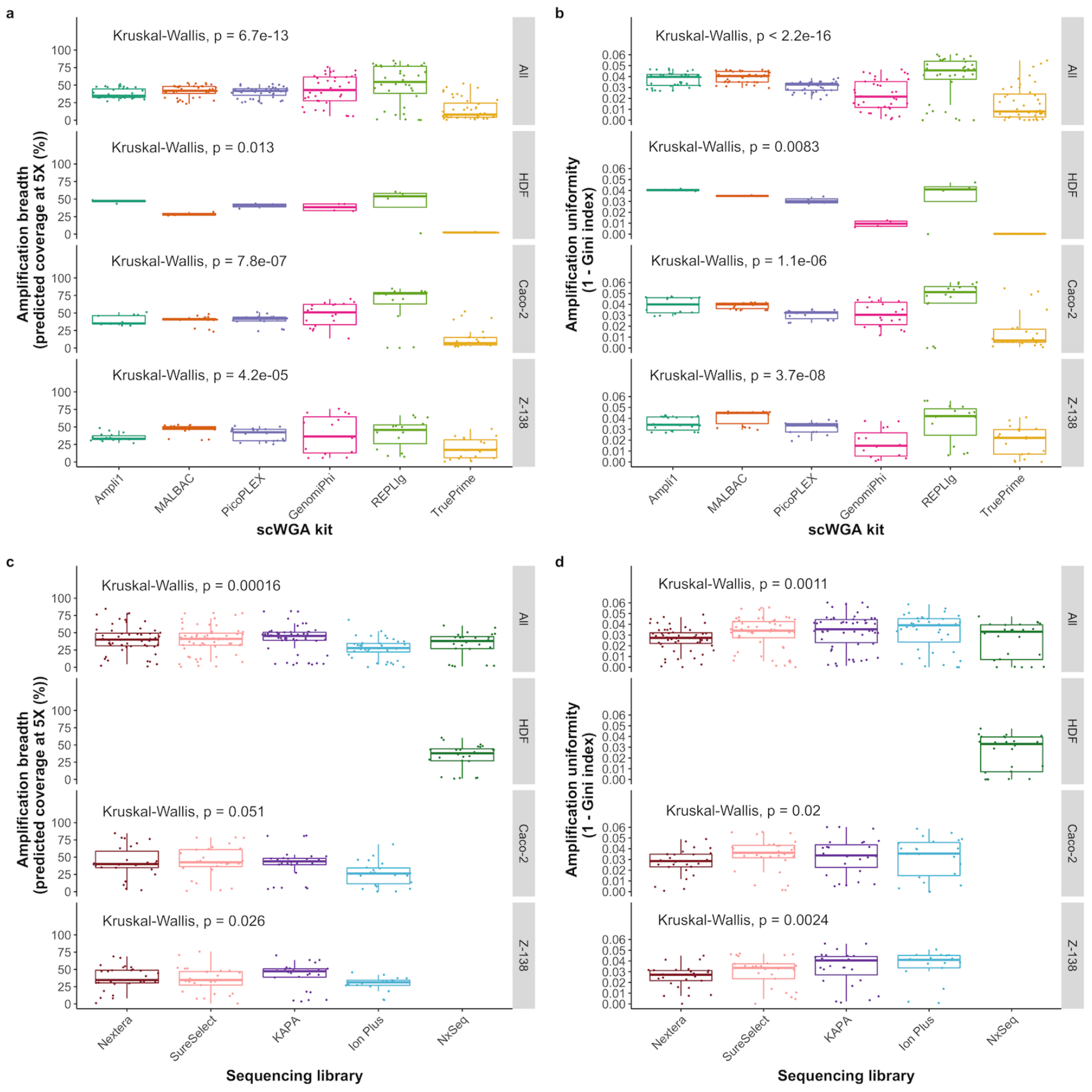
Amplification breadth and uniformity. **a,b,** Effect of the scWGA method on (**a**) predicted amplification breadth, measured by the percentage of the genome predicted to be covered at 5X, and (**b**) amplification uniformity, measured as 1 minus the Gini index of the resulting Lorenz curves, for the different scWGA kits. **c,d,** Effect of the sequencing library on (**c**) predicted amplification breadth and (**d**) amplification uniformity. **a-d,** Boxplots: the central line indicates the median, while the box limits correspond to the Q1 and Q3 quartiles; upper and lower whiskers extend from Q3 to Q3 + 1.5 × (Q3 - Q1) and from Q1 to Q1 - 1.5 × (Q3-Q1), respectively. Kruskal-Wallis test p-values are shown.

Albeit less dramatic, the library protocol also had a significant effect on amplification breadth and uniformity. Our modified KAPA protocol provided more breadth than the other library protocols, with Ion Plus being the worst in this aspect (Fig. 4c). Also, Ion Plus and Nextera yielded overall the most and least uniform amplifications, respectively (Fig. 4d). As expected, the two sequencing technologies used, Illumina and Ion Torrent, did not have a significant effect on the uniformity of the amplification. In order to better understand the joint effect of the different parameters (cell line, scWGA kit, amplification location, library kit, yield DNA, amplicon size, sequencing depth and sequencing technology) on amplification uniformity we fitted a multivariable regression model upon the Gini index values, finding that differences in amplification uniformity could be explained by the amplification kit alone.

### Amplification recurrence

The coverage distribution along the genome observed for the single-cells was significantly correlated with that of the unamplified bulk (Fig. 5). Importantly, we also found that two cells amplified with the same scWGA kit showed significantly more regions in common than two cells amplified with a different scWGA kit, except for REPLIg (Fig. 5 and Supplementary Figs. 2-4). In addition, we observed that non-MDA showed a significantly higher coverage in regions with high GC content, as previously reported^31^. Interestingly, REPLIg showed a negative correlation of coverage with GC content whereas the other MDA methods did not show any preference based on sequence content.

**Fig. 5.**
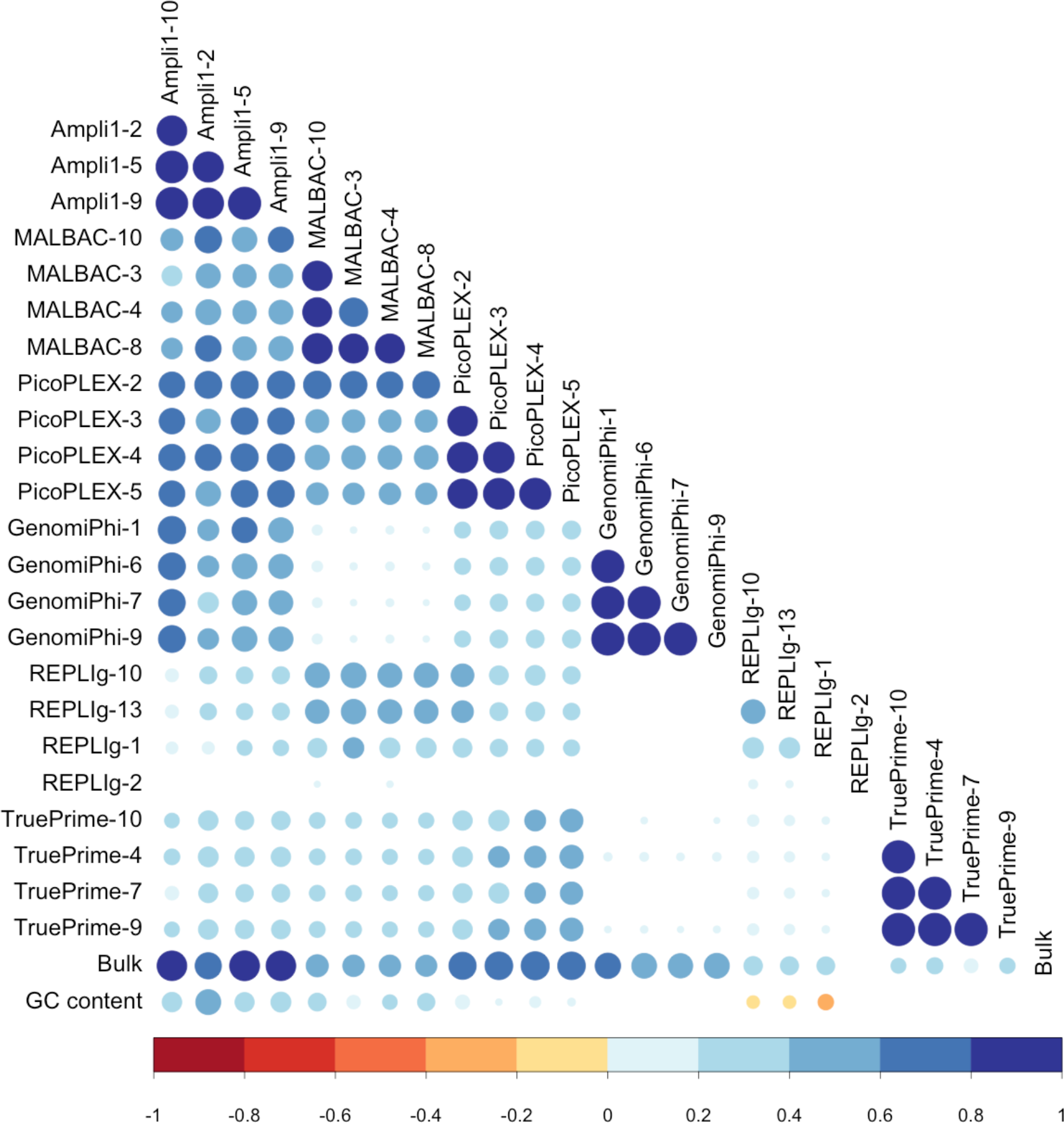
Amplification recurrence. Correlation plot of read counts along 1 Mb-long genome windows. The size of the dots indicates the Pearson correlation coefficient value from low (small) to high (big) and colored differently if positive (blue scale) or negative (red scale). Only statistically significant values (p-value < 0.05) are shown.

### Chimera rates, allelic imbalance, ADO and false SNVs

During scWGA, several artifacts can be produced, such as the formation of chimeric molecules, biased amplification of alleles, and amplification errors. These errors can easily result in incorrect genotype calls. Here we measured chimera rates, allelic imbalance, allelic dropout (ADO) and false positive variant calls (Fig. 6 and Supplementary Table 7). In this case, only the HDF cell line (4 cells per scWGA method) was used as it lacks somatic variation, which otherwise could have easily confounded these estimates. Chimera rates were much higher (>10%) for GenomiPhi and TruePrime, with Ampli1 and MALBAC showing the lowest values (Fig. 6a). When these rates were estimated upon paired-end discordant reads instead of split reads, the trends were the same but the rates were twice as high (Fig. 6b).

**Fig. 6.**
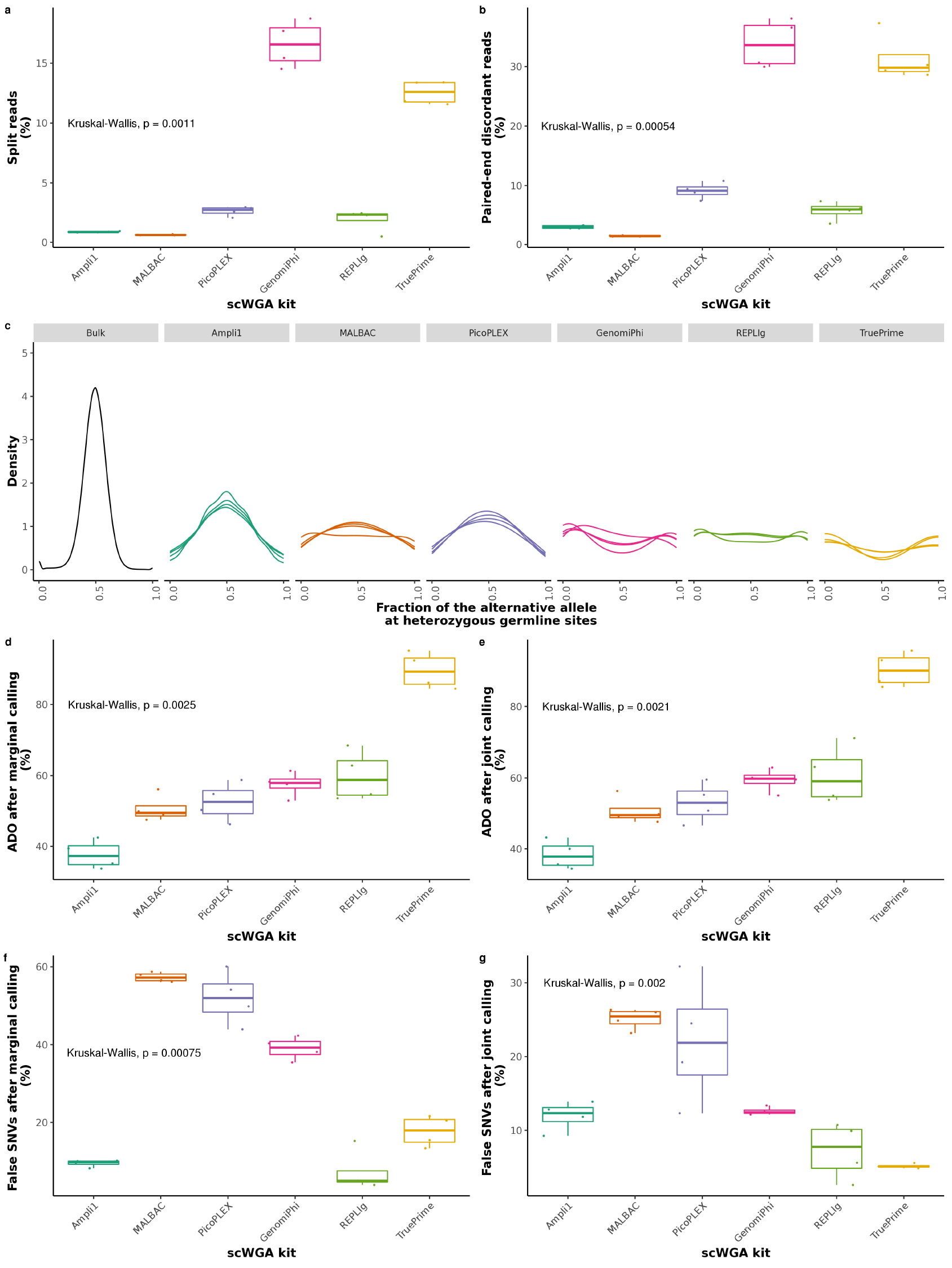
Chimera rates, allelic imbalance, ADO and false SNV calls. **a-g**, all these values were measured from the healthy cell line HDF. **a**, paired-end discordant reads. **b,** split reads. **c**, Kernel density estimation of the alternative allele fraction at heterozygous germline sites. Each line within a panel tab represents a different single-cell. For the density estimation we used all the positions called as heterozygous in the bulk with *HaplotypeCaller* and with at least 6 reads of coverage in the single-cells. The density of one of the REPLIg single-cell is not shown as it only had 7 positions with more than 5 reads, too few to obtain a smooth distribution of the allele fraction. **d**, ADO calculated after marginal genotype calling. **e**, ADO calculated after joint genotype calling. **f,** False SNVs calculated after marginal genotype calling. **g,** False SNVs calculated after joint genotype calling. **d-g,** Boxplots: the central line indicates the median, while the box limits correspond to the Q1 and Q3 quartiles; upper and lower whiskers extend from Q3 to Q3 + 1.5 × (Q3 - Q1) and from Q1 to Q1 - 1.5 × (Q3-Q1), respectively.

In terms of allelic imbalance and ADO rates, non-MDA methods outperformed MDA methods (Fig. 6c-e). Still, ADO rates were very high in all cases, ranging from the 38% rate of Ampli1 to the 60-90% values observed for TruePrime. REPLIg, Ampli1, and TruePrime resulted in more accurate SNV calls than the other methods (Fig. 6f,g), although the false positive rates obtained were halved depending on the variant calling approach (marginal vs joint; see methods). False SNVs were usually biased towards transitions, but different scWGA approaches showed somehow different error signatures (Supplementary Fig. 5). In the case of Ampli1, these errors showed a pattern most similar to the human germline mutational profile, in which transitions occur more than twice as often as transversions. Besides, we also found an increase in G:C>A:T errors in REPLIg and GenomiPhi.

### Copy-number detection

Copy number aberrations are a fundamental type of structural variation of interest to assess genomic heterogeneity among cells. The copy-number profiles estimated from the diploid HDF cell line were generally accurate (i.e., we expect a copy number of 2 for all genomic regions) for all scWGA methods, except for TruePrime (Supplementary Fig. 6d-i). The coverage dispersion measure (MAD)^32^ was significantly smaller for non-MDA methods, although for Caco-2 REPLIg showed also very low MAD values (Supplementary Fig. 6a-c).

### Mapping rates and duplicated reads

Finally, we explored the mapping rates and percentage of duplicated reads obtained. For all scWGA methods, we observed a high percentage of mapped reads (Supplementary Tables 4-6), with marginally significant differences among them (*p-value* = 0.04). In particular, TruePrime showed more reads mapped to the mitochondrial genome than the other scWGA kits (close to 9% in Caco-2, up to 6% in Z-138, and as high as 84.71% in HDF). Also, we detected clear mapping differences among the sequencing library kits (*p-value* < 2.2e-16). Within these, the library protocols that include an enrichment PCR step (SureSelect and Nextera) showed a significantly higher percentage of mapped reads (*p-value* = 3.8e-15). Additionally, we found a clear effect of the sequencing technology on the percentage of duplicates estimated, with Ion Torrent producing significantly more duplicates than Illumina (24% and 6%, respectively; *p-value* < 2.2e-16).

### Benchmark of the breadth and uniformity measures

As an estimate of amplification breadth, we used low-pass sequencing data to predict the fraction of the genome covered at higher sequencing depths with the package Preseq^33^. To evaluate amplification uniformity, we used the Gini index or area below the coverage Lorenz curve. To validate both measurements we leveraged an *in-house* dataset of 30 single cells from a Chronic Lymphocytic Leukemia (CLL) patient. The analysis of these data demonstrated that the amplification breadth prediction was very accurate (Supplementary Fig. 7). In addition, both the breadth prediction and the 1 – Gini index showed a significant positive correlation with the percentage of chromosomes amplified (Supplementary Fig. 8).

## Discussion

Overall, our results show that MDA approaches (GenomiPhi, REPLIg, TruePrime) produced higher yields than non-MDA methods (Ampli1, MALBAC, PicoPLEX), which could be related to a more stable polymerase activity under isothermal conditions^34^. At the same time, MDA approaches generated larger amplicon sizes than non-MDA methods, likely due to a higher processing capability and template affinity of the Phi29 polymerase^35^. In particular, REPLIg clearly outperformed the other scWGA strategies in this aspect, possibly resulting from a higher DNA polymerase concentration^36^.

TruePrime resulted in a significantly lower coverage breadth, as previously reported^28^, with REPLIg outperforming the remaining methods in general, but with higher variance, as the other MDA methods. This might be explained, at least partially, by the random annealing of primers and/or more variable amplicon sizes of the MDA methods. In agreement with the coverage breadth predictions, REPLIg and TruePrime resulted in the most and least uniform amplifications, respectively, measured by the variation of the coverage along the genome. The lysis and denaturalization steps in TruePrime are made on ice, which might prevent the availability of single-stranded DNA for the primase. On the other hand, REPLIg has already been suggested to provide a more uniform coverage than non-MDA methods for few-cell amplification^30^.

The observed coverage correlation along the genome between single cells and the unamplified bulk was expected, mainly due to mappability limitations for repetitive sequences^37^. However, unless there is a recurrent amplification bias, we would not expect to see higher correlations among cells amplified with the same scWGA kit compared to cells amplified with a different scWGA kit, or compared with the bulk. Therefore, our results suggest that the amplification bias along the genome is not random for the different scWGA kits studied, except for REPLIg, which in most cases resulted in apparently random amplification. For non-MDA methods, spatial recurrence in the amplification is expected because they use their own set of non-random primers. For MDA methods, the results are more difficult to interpret. TruePrime uses a primase that generates its own primers^9^, while for both GenomiPhi and REPLIg random primers are added. However, only REPLIg showed a random amplification bias. Zhang et al.^12^ obtained a better fit with a statistical model with random amplification bias for MDA, but they do not clarify which exact MDA method they used.

For the HDF cell line, non-MDA methods showed a distribution more similar to the bulk for the alternative allele frequency in heterozygous germline sites, clearly outperforming the MDA methods in terms of allelic imbalance. In particular, Ampli1 seemed to produce very low allelic imbalance. This good behavior of Ampli1 might be the result of a synthesis procedure that converts residual single-strand DNA (ssDNA) molecules into double-strand DNA (dsDNA) molecules. Also, in PicoPLEX and MALBAC the amplification is quasi-linear, therefore limiting the propagation of any allelic bias. We would like to remark that, in general, alternative allele frequency distributions are wider for our single cells than for the bulk not only because of allelic imbalance but also because they have lower coverage at the sites considered.

In agreement with these results –ADO is an extreme case of allelic imbalance–, non-MDA methods showed much lower ADO rates than MDA methods, in particular Ampli1, who showed the lowest ADO rate (< 40%). On the other extreme, TruePrime showed a large ADO rate (> 80%). Still, a 40% ADO rate is much higher than previously reported for Ampli1^25,38,39^. Although values over 40% have been already observed in other experiments for MDA methods^40^, here the ADO rates for GenomiPhi and REPLIg were closer to 60%. Admittedly, it is possible that the absolute ADO values estimated here are somehow inflated due to the small coverage. It is well-known that at sites with low coverage there is a higher probability of missing one of the alleles by chance^41^. Although we have carried out this analysis on sites with at least 6X, we are aware that at this depth it is possible to incorrectly call as homozygous a truly heterozygous site. Indeed, genotyping of heterozygous sites only approximates a correct call rate of 1.0 for coverages higher than 15X^42^, although a threshold value of 6X has been used before to estimate ADO and false SNV rate^40^. In any case, the relative ADO performance should still correspond with the trend observed.

We used false SNVs as proxies for amplification errors. Indeed, the former also include sequencing errors, wrong SNV calls and potentially some true somatic variants, so the absolute value might be more or less inflated. However, the comparison of the false SNVs observed for each scWGA method should inform us about their relative amplification error rates. We used a PCR-free library protocol, and sequencing errors should be similar for all 24 HDF single-cell libraries, as they were included in the same sequencing run. Also, most of the true somatic variants should have been filtered out with the help of Monovar. Finally, whatever it is, the number of calling errors should be more or less constant across cells. REPLIg and TruePrime resulted in the lowest false SNV rates (< 8% depending on the genotyping approach). This might be related to the polymerase used for amplification, Phi29, which has a much lower error rate than the ones used by the other scWGA kits^8,10,11,14,43–45^. REPLIg was already reported as having low error rates^17,28^. Ampli1 also performed quite well in this regard, perhaps due to the use of a combination of Taq with a proofreading polymerase Pwo^7^ with low error rates^46^. Importantly, when we used joint variant calling –incorporating population-level information–, the inferred false positive rates were halved. With regard to the type of errors observed, we found that both REPLIg and GenomiPhi resulted in an excess of C:G>T:A transitions, which has been previously attributed to high-temperature denaturation protocols, whereas the signature for Ampli1 better reflects the one expected for unamplified bulk samples^47^. Being aware of the existence of a different error signature for the different scWGA kits is a fact to consider if one is interested in detecting mutational signatures in single cells.

Our results suggest that MALBAC and Ampli1 form fewer chimeras than the other methods. Indeed, chimeras can bias the inference of structural variants. It was expected for MDA methods to perform worse than non-MDA methods in this regard, due to the Phi29 DNA polymerase strand displacement activity resulting in chimeric molecules^18^. However, despite the fact that REPLIg is based on MDA, it showed very low chimera rates, even lower than microfluidic protocols^48^. Perhaps, this might be related to larger amounts of DNA polymerase which could limit the time that ssDNA strands are naked and available for chimera formation.

For the HDF cell line, copy-number profiles were in general accurate for all scWGA methods except for TruePrime. In the other cell lines, the copy-number profiles were much more segmented. Nevertheless, read counts were much more dispersed for MDA methods (except for REPLIg in the HDF cell line), which could be partially explained by uneven amplification (Supplementary Fig. 6a-c). Consequently, non-MDA methods would be the recommended choice for CNV analysis, as suggested before^11,38^.

Not surprisingly, the amplification protocols did not significantly affect the mapping rates. While the scWGA kit employed had a much smaller effect on the percentage of mapped reads than the library construction method –strategies with an enrichment PCR step like SureSelect and Nextera were better–, in all cases these percentages were quite high. As expected, probably due to the emulsion PCR step^49^ included in the Ion Torrent sequencing protocol, the latter showed significantly more duplicates than Illumina.

In conclusion, none of the scWGA methods outperformed the others in all scenarios assessed, but clearly, some are better than others in different aspects. Here we have exposed distinct advantages and weaknesses of different scWGA methods that will be important for the interpretation and analysis of single-cell genomes.

## Methods

### Cell-lines

We used three different cell lines for the different experiments, HDF, Caco-2, and Z-138. HDF is a healthy neonatal, diploid fibroblast cell line (HDF) purchased from Sigma-Aldrich (https://www.sigmaaldrich.com). Caco-2 is a polyploid colorectal cancer cell line with a modal chromosome number of 96 purchased from the American Type Culture Collection (ATCC; https://www.atcc.org). The Z-138 is a hyperdiploid mantle cell lymphoma cell line with a modal chromosomal number of 49, also purchased from ATCC. We cultured all cell lines under an atmosphere containing 5% CO_2_ at 37°C. We grew HDF in an all-in-one ready-to-use fibroblast growth media (Sigma-Aldrich), Caco-2 in a media consisting of Dulbecco’s Modified Eagle’s Medium/F12 with 3.151 g/l glucose and L-glutamine (Lonza) and Z-138 with Iscove’s Modified Dulbecco’s Medium (ATCC). For Caco-2 and Z-138, we completed the media with 10% fetal bovine serum EU standard (Biochrom) and 1% penicillin/streptomycin (Lonza) at a working concentration of 100 units of potassium penicillin and 100 μg of streptomycin sulfate per 1 ml of culture media.

### Single-cell isolation

We cultured HDF and Caco-2 cells until 80% confluence before staining with Hoechst 33342 (BD Biosciences) following the fabricant’s recommendations. Thereafter, we harvested, resuspended at a concentration of 10^6^ cells per ml in phosphate buffered saline (PBS) and marked cells with propidium iodide (PI; BD Pharmingen). Then, we sorted live single cells in G0/G1 with a BD Biosciences FACSAria III flow cytometer (BD Biosciences, Madrid, Spain) and collected them into 96 well plates with 1-3 μl of PBS (Supplementary Fig. 9). For sorting, we used the BD FACSDiva v8.0.1 (BD Biosciences, Madrid, Spain) and FlowJo v7.6.2 (FlowJo, LLC, Ashland, OR, USA) for further analyses. For Z-138 we followed the same strategy but without Hoechst staining and using a FACS Aria 2.0 (BD Biosciences, Madrid, Spain) for sorting. Single cells were stored at −80 °C until amplification.

### Single-cell whole-genome amplification

We used six different kits for single-cell whole-genome amplification (scWGA): Ampli1 (Silicon Biosystems), Multiple Annealing and Looping Based Amplification Cycles (MALBAC; Yukon Genomics), PicoPLEX (Rubicon Genomics), Illustra Single Cell GenomiPhi (GE Healthcare), REPLIg Single-Cell (Qiagen) and TruePrime (SYGNIS) following the manufacturer’s protocols (Supplementary Table 2). In order to reduce contamination, we carried out scWGA in a laminar-flow hood using a dedicated set of pipettes and UV irradiated plastic materials. We also included positive (10 ng/μl REPLIg human control kit, QIAGEN) and negative (DNase/RNase free water) controls. For Ampli1 we carried a few extra steps after amplification. We used the Ampli1 QC kit to select those amplification products that were positive for four PCR markers. In order to increase the total dsDNA content, we used the Ampli1 ReAmp/ds kit. Afterwards, we removed the adaptors adding 5 μl of NEBuffer 4 10X (New England Biolabs), 1 μl of MseI 50U/μl (New England Biolabs) and 19 μl of nuclease-free water to 25 μl of dsDNA, using a thermal cycler at 37 °C for 3 h, followed by enzyme inactivation at 65 °C for 20 min.

Attending to the fabricant’s recommendations, we purified PicoPLEX and MALBAC products with the QIAquick PCR Purification protocol (Qiagen) and Ampli1 products with 1.8X AMPure XP beads (Agencourt, Beckman Coulter). MDA methods do not include a purification step. We measured DNA yield with a Qubit 3.0 (ThermoFisher Scientific) fluorometer and amplicon fragment size with a 2200 TapeStation platform (Agilent Technologies). We measured amplicons from non-MDA based scWGA methods using the D5000 ScreenTape System and amplicons from MDA-based kits using the Genomic DNA ScreenTape System. The latter also allowed us to measure the integrity of the amplicons (DNA Integrity Number or DIN).

In total, we amplified 230 single-cells: 34 with Ampli1 (4 HDF, 12 Caco-2 and 18 Z-138), 40 with MALBAC (4 HDF, 18 Caco-2 and 18 Z-138), 40 with PicoPLEX (4 HDF, 18 Caco-2 and 18 Z-138), 37 with GenomiPhi (4 HDF, 18 Caco-2 and 15 Z-138), 35 with REPLIg (4 HDF, 16 Caco-2 and 15 Z-138) and 44 with TruePrime (4 HDF, 22 Caco-2 and 18 Z-138) (Fig. 2 and Supplementary Table 2).

### Bulk DNA extraction

We extracted bulk genomic DNA (gDNA) from the HDF cell line with the QIAamp DNA Mini kit (QIAGEN) according to the fabricant’s recommendations. We estimated concentration and gDNA integrity as previously described for single cells.

### Next-generation sequencing libraries

We built 230 single-cell whole-genome libraries employing five different library preparation kits: SureSelect^QXT^ (Agilent Technologies), NxSeq AmpFREE Low DNA (Lucigen), Ion Plus Fragment library (ThermoFisher Scientific), Nextera DNA (Illumina) and KAPA (Kapa Biosystems). We built SureSelect and NxSeq libraries following the commercial indications while we slightly modified Ion Plus and Nextera. The sequencing facility of the National Center for Genomic Analysis (CNAG; http://www.cnag.crg.eu) modified the KAPA protocol to a larger extent (Supplementary Table 3). We mechanically sheared the DNA in an S2 or a LE220 Focused-ultrasonicator (Covaris) for the NxSeq, Ion Plus and KAPA protocols (Supplementary Table 8), while for SureSelect and Nextera we fragmented the DNA enzymatically. For Ion Plus, we included an extra purification step with AMPure XP beads (1.2X beads/sample ratio) (Agencourt, Beckman Coulter) for a better removal of NGS adaptors. For Nextera, we used 200 μl of washing buffer instead of the 300 μl recommended by the provider, the centrifugation speed was 10,000 g at room temperature (RT) for 30 s instead of 1,300 g at 20 °C for 2 min, and we used AMPure XP beads in a 0.8X ratio instead of 0.6X. In addition, we eluted the Nextera libraries in 20 μl of resuspension buffer instead of 32.5 after a 5 min air-dried step instead of 15. For the modified KAPA protocol, we carried out the end-repaired of 500 ng of sheared DNA, adenylation and ligation to Illumina specific indexed paired-end adaptors (NEXTflex-96^™^ DNA Barcodes, Bio Scientific). We performed the DNA size selection in two steps (0.65X and 0.85X beads/sample ratio) with AMPure XP beads in order to reach the desired fragment size (450 bp). Finally, we measured library insert sizes with a 2100 Bioanalyzer High Sensitivity DNA Kit (Illumina) or a 2200 TapeStation High Sensitivity D1000 (Ion Torrent). We quantified library concentration with the Kapa Library Quantification Kit (Kapa Biosystems) for Illumina and the Ion Library Quantitation Kit (Life Technologies) for Ion Torrent.

In addition, we constructed a whole-genome HDF bulk library using NxSeq AmpFREE Low DNA (Lucigen) (Supplementary Table 3). We performed size selection using AMPure XP beads and we checked fragments size with the 2100 Bioanalyzer High Sensitivity DNA Kit. Finally, we quantified library concentration using the KAPA Library Quantification Kit.

### Whole-genome sequencing

We sequenced single-cell libraries at shallow depths (0.07-1.76X). We sequenced 24 HDF and 166 Caco-2/Z-138 libraries on an Illumina HiSeq 4000 (PE150) or HiSeq 2000 (PE125), respectively, at CNAG. We sequenced the remaining 40 libraries with an Ion Proton platform (Ion PI chip v3) at the Galician Public Foundation of Genomic Medicine (FPGMX; http://www.xenomica.eu). We sequenced the bulk library of HDF at 30X on an Illumina HiSeq 4000 (PE150) at CNAG.

### Preprocessing of NGS data

We clipped library adapters and also those included in the Ampli1, PicoPLEX and MALBAC amplifications using CutAdapt (v.1.11,v.1.14)^50^. We mapped the sequencing reads with at least 70 bp to the human reference genome (hs37d5) with BWA-MEM (v.0.7.15-r1140)^51^. We sorted reads and flagged duplicates with Picard *SortSam* (v.2.2.1; http://broadinstitute.github.io/picard) and Picard *MarkDuplicates*, respectively. We independently mapped reads from different lanes and then merged during the duplicate marking process taking into account their read group. Regarding Ion Plus libraries, the sequencer software already removed the adapters. In order to map the reads with the Torrent Mapping Alignment Program (TMAP; v.3.4.1; https://github.com/iontorrent/TMAP) to the hs37d5 genome, we had to transform first of all the original BAM files to FASTQ format using Picard *SamToFastq*. We also sorted the BAM files and marked duplicates as explained above.

For the 24 HDF single-cells and bulk, which were subsequently used for variant calling, the base quality scores were recalibrated for each sample using GATK (v.3.7)^52^. Afterward, we realigned reads from the single-cells and bulk together around known indels to avoid potential false SNV calls not related to the amplification process itself but due to misalignments.

### Amplification breadth

In order to approximate the amplification breadth, we calculated the percentage of the genome that would be covered by one or more reads (coverage breadth). However, we did not calculate this value directly because at very low depths this estimate is not reliable due to sampling error^12^. Instead, we used a method developed for single-cell libraries, implemented in the *gc_extrap* function of the package Preseq^33^. For this calculation, we downsampled the BAM files to the lowest depth observed (0.07X) with Picard *DownsampleSam.* To check the quality of the Preseq prediction, we took advantage of two *in-house* datasets consisting of 30 single-cells from a patient with Chronic Lymphocytic Leukemia (CLL), that were amplified with REPLIg and sequenced twice at 0.3X-0.6X and 5X. We downsampled the former to exactly 0.1X and predicted with Preseq the percentage of the genome covered by one or more reads at 5X. Then, we compared this prediction with the values observed for the 5X dataset. In addition, we compared the Preseq prediction and the Gini index (see below) with the percentage of chromosomes amplified in the same 30 CLL single cells, according to a panel of 20 PCR markers representing all human chromosomes except 6 and 11.

### Amplification uniformity

We used coverage uniformity as a surrogate for amplification uniformity. For this, we used the downsampled BAM files at 0.07X. We calculated the sequencing depth per site with Bedtools^53^ and parsed its output with an *in-house* script to create the Lorenz curves^47^ with the *Lc* function of the *Ineq* R package. In order to quantitatively compare the Lorenz curves, we calculated the Gini index for each cell^20^. The Gini index measures the area below the Lorenz curve, spanning between 0 and 1, being 0 perfect uniformity and 1 perfect disuniformity, so we defined amplification uniformity as 1 minus the Gini index (see also Supplementary Fig. 1). We estimated the Gini indexes using the *Gini* function from the *Ineq* package. In order to understand which variables of the study (cell line, scWGA kit, amplification location, library kit, DNA yield, amplicon size, sequencing depth and sequencing technology) affect most the Gini index, we fitted a regression model.

### Amplification recurrence

In order to understand whether a given scWGA method tends to preferentially amplify the same genomic regions in different cells compared to other scWGA methods, we counted the number of reads within non-overlapping windows of 1 Mb with Pysamstats (https://github.com/alimanfoo/pysamstats). For this, we used downsampled BAM files at 0.1X –after removing duplicates, secondary alignments and unmapped reads with Samtools *view^54^*– as input. We only used the HDF cell line for this calculation in order to avoid potential correlations among read counts due to a heterogeneous copy number in the cancer cell lines. Subsequently, we computed the Pearson correlation coefficient of the read counts between pairs of single-cells (amplified with the same scWGA kit or not) and between single-cell and bulk. We further explored the correlations between read counts and GC content in sliding windows. GC content was assessed with Bedtools *nuc*. However, since having for instance 2 reads of 150 bp mapped to the same 1 Mb window in two different single-cells does not necessarily mean that they amplify exactly the same region (in one cell reads could be mapped to the first 500 bp of the window and in the other cell to the last 500), we further explored presence/absence amplification recurrences computing Jaccard similarity coefficients. For this, we created bedGraph files for each of the single cells using Bedtools *genomecov*, simplified integer coverage values simply to 1 (presence) or 0 (absence) and then merged the resulting files using Bedtools *unionbedg* to get a matrix. We removed consecutive matrix rows which showed exactly the same presence/absence profile for the 24 HDF cells, as we wanted to be stringent and do not count several events from a single read recurrence. Then, we computed the Jaccard similarity coefficient for each pair of single-cells using the R package *jaccard* (https://cran.r-project.org/web/packages/jaccard/index.html). Basically, the Jaccard coefficient is computed by dividing the number of intersections (presence of coverage in both cells) between the number of total unions (presence of coverage in just one cell), this way ignoring positions without coverage for the two cells (avoiding increasing the similarity due to random absence of coverage originated by low pass sequencing). To assess its statistical significance, we implemented a permutation test using a homemade R script.

### Chimera formation rate

We considered the paired-end reads mapping at a distance higher than 1 kb to result from chimeric amplicons, as well as reads with supplementary alignments (split reads). We calculated paired-end distances using Picard *CollectAlignmentSummaryMetric* and identified split reads through detection of SA:Z tags in the BAM files. For this calculation, we only used the HDF cell line in order to avoid false positives due to the high genetic instability in Caco-2 and Z-138.

### Allelic imbalance and ADO

During scWGA, the two alleles of a diploid single-cell can be amplified in an unequal manner. Deviations of allele frequencies at heterozygous germline sites evince such events. If these frequencies are different from the theoretical 50%, we consider that an allelic imbalance event, and if the deviation is so high that one of the alleles is completely lost and it cannot be detected, we designate it as ADO (Fig. 1). Again, for these calculations, we only used the HDF cells, as Caco-2 and Z-138 cells present variable ploidy.

#### Allelic imbalance

We ran GATK *HaplotypeCaller* for the HDF bulk with the parameter--pcr_indel_model set to NONE. We used GATK *SelectVariants* to keep the heterozygous sites and ran GATK *VariantRecalibration* (v. 4.0.0.0) to select a high confidence set. In parallel, we created pileup files from all the HDF single-cells and bulk with Samtools *mpileup* and extracted the alternative allele fraction at the high confidence heterozygous positions using a Python script (Supplementary Information). We only considered the allele frequencies derived from sites covered by at least six reads. In order to obtain a smooth distribution from the discrete counts, we estimated the probability density function of the alternative allele fractions with the *core* R stats package, with the *bandwidth adjust* parameter set to three.

#### Allelic dropout (ADO)

In order to measure ADO, we first had to obtain genotypes both for the bulk and the single cells. For this, we ran GATK *HaplotypeCaller* in ERC mode for all HDF single-cells and bulk independently (for the latter again setting--pcr_indel_model to NONE) and merged the results using two different approaches with GATK *GenotypeGVCF*. On one hand, we aggregated the bulk and the 24 single-cells (“*joint calling*”) and on the other hand, we simply combined one single-cell and the bulk one at a time (“*marginal calling”*). We counted an ADO event when the bulk was genotyped as heterozygous (AB) and the single-cell as homozygous for either the reference or the alternative allele (AA or BB) (Supplementary Fig. 10). For this calculation, we only considered positions covered by six or more reads in single cells and 15 or more reads in the bulk.

### Amplification polymerase errors

We used false SNV calls (“false positives” or FP) as proxies for amplification errors of the polymerase. Using the same genotypes estimated above for the ADO calculation, we counted an FP event when the bulk genotype was homozygous (AA or BB) and the single cell heterozygous (AB) (Supplementary Fig. 10). However, given the high number of somatic mutations expected to accumulate during cell growth on a plate and with the intention of mitigating their effect on the calculation of the FP rate, we did not consider for this calculation SNVs detected by the single-cell variant caller MonoVar^55^, nor sites with two or more reads containing the “potentially erroneous allele” in the bulk. That is, we preferred not to be very stringent and allowed one error in the bulk potentially arising by a sequencing or mapping error. Again, we only considered positions with six or more reads for single cells and 15 or more reads for the bulk. We also explored the mutational profile (signature) of the FPs directly extracting alternative and reference alleles from the marginal calling VCFs and grouped them into different categories.

### Copy-number detection

The uneven distribution of the coverage, added to the existence of ADO, can make copy number detection from single-cell problematic. We compared how different scWGA kits behave in this regard using the coverage dispersion measure (MAD) explained in Garvin et al.^32^. For this calculation, we used the downsampled BAM files at the lowest depth for each cell line (0.25X, 0.07X, and 0.08X for HDF, Caco-2 and Z-138, respectively). We filtered out reads with a mapping quality lower than 20 from the downsampled BAMs using Samtools and created with Bedtools the BED files required to run Ginkgo^32^. We ran Ginkgo under default settings except for the Binning Simulation Options that were set to 150 bp reads and BWA mapping. The segmentation was inferred independently for each sample to allow a fair comparison.

### Statistical tests

We calculated descriptive statistics using the *kruskal.test*, *wilcox.test*, and *cor.test* functions available in the *core* R stats package^56^. For the regression models, we used the R package *FWDselect*^57^.

## Code availability

The R and bash scripts used for this study are available as Supplementary Information.

## Data availability

Bulk and single-cell FASTQ files generated for this study have been deposited at the Sequence Read Archive (SRA) under STUDY accession SRP162960.

## Acknowledgments

This work was supported by the European Research Council (ERC-617457- PHYLOCANCER awarded to D.P.) and by the Spanish Ministry of Economy and Competitiveness - MINECO (BFU2015-63774-P awarded to D.P.). D.P. receives further support from Xunta de Galicia. N.E.-G. is supported by a Ph.D. fellowship from the Galician Government (ED481A-2015/475). T.P. is now supported by a Ph.D. fellowship from the Spanish Government (FPU15/03709) and previously by a Ph.D. fellowship from the Galician Government (ED481A-2015/083). H.H. is a Miguel Servet (CP14/00229) researcher funded by the Spanish Institute of Health Carlos III (ISCIII). H.H. receives funding from the Ministerio de Ciencia, Innovación y Universidades (SAF2017-89109-P; AEI/FEDER, UE). Core funding is from the ISCIII and the Generalitat de Catalunya. CLL samples included in this study were provided by the IISGS Biobank (PT13/0010/0022), integrated into the Spanish National Biobank Network and they were processed following standard operating procedures with the appropriate approval of the Ethical and Scientific Committees. We want to thank Pilar Alvariño and people from FPGMX for their help with some of the experiments, and Laura Tomás, Fabián Crespo, Harald Detering, Xose S. Puente, Andrés Pérez-Figueroa and Joao Alves for discussions. We thank the Supercomputation Center of Galicia (CESGA) for providing computational resources.

## Author contributions

N.E.-G., S.P.-L., A.G.-A. and H.H. performed the experiments. T.P. carried out the bioinformatic analyses. N.E.-G. and T.P. performed the statistical analyses. N.E.-G., T.P., S.P., and D.P. wrote the manuscript. All authors read, commented and approved the final manuscript. D.P. designed the project and supervised the whole study.

